# Image segmentation and separation of spectrally similar dyes in fluorescence microscopy by dynamic mode decomposition of photobleaching kinetics

**DOI:** 10.1101/2022.02.28.482234

**Authors:** Daniel Wüstner

## Abstract

**Background:** Image segmentation in fluorescence microscopy is often based on spectral separation of fluorescent probes (color-based segmentation) or on significant intensity differences in individual image regions (intensity-based segmentation). These approaches fail, if dye fluorescence shows large spectral overlap with other employed probes or with strong cellular autofluorescence.

**Results:** Here, a novel model-free approach is presented which determines bleaching kinetics based on dynamic mode decomposition (DMD) and uses the inferred photobleaching kinetics to distinguish different probes or dye molecules from autofluorescence. DMD is a data-driven computational method for detecting and quantifying dynamic events in complex spatiotemporal data. Here, DMD is used to determine photobleaching characteristics of a fluorescent sterol probe, dehydroergosterol (DHE), compared to that of cellular autofluorescence in the nematode Caenorhabditis elegans. It is shown that decomposition of those dynamic modes allows for precise image segmentation, thereby separating probe from autofluorescence without invoking a particular model for the bleaching process. In a second application, DMD of dye-specific photobleaching is used to separate two green-fluorescent dyes, an NBD-tagged sphingolipid and Alexa488-transferrin, thereby assigning them to different cellular compartments.

**Conclusions:** Data-based decomposition of dynamic modes can be employed to analyze spatially varying photobleaching of fluorescent probes in cells and tissues for image segmentation, discrimination of probe from autofluorescence and image denoising. The new method should find wide application in analysis of dynamic fluorescence imaging data.

## Background

Image segmentation in fluorescence microscopy is either based on differences in intensity of one fluorescent probe, e.g., DAPI intensity is selective for the nucleus but not found in other organelles, or on using specific dyes for each subcellular compartment. Selective detection of different fluorescent molecules in intracellular organelles requires differences in emission wavelength (color) but this approach fails if spectral properties of dyes are very similar. Spectral unmixing can overcome this limitation to some extend but only, as long as the excitation or emission peaks of individual fluorophores differ by at least 30 nm [1]. Dye-specific properties, such as fluorescence lifetime can also be employed for selective separation of different probes or of probe from autofluorescence [2], but this requires special equipment not being available in many cell biological laboratories.

Photobleaching kinetics of, for example organelle markers, can also be used to segment intracellular organelles or to distinguish probe intensity from autofluorescence [3]. This is particularly important, if excitation and fluorescence spectra of a probe and of autofluorescence overlap strongly and can hardly be distinguished based on intensity differences. One such application is the analysis of sterol trafficking in the nematode Caenorhabditis elegans (C. elegans). C. elegans is a sterol-auxotroph organism, which is often employed to study the molecular basis of lipid transport and metabolic regulation on a systemic level [4-6]. Almost the entire sterol pool of this nematode can be replaced by feeding these worms with the fluorescent natural sterol dehydroergosterol (DHE), allowing for observing sterol uptake and transport by microscopy [7, 8]. To detect DHE selectively and to distinguish it from cellular autofluorescence in the ultraviolet, we made use of the much faster photobleaching of the sterol compared to autofluorescence of subcellular structures, particularly of gut granules [8, 9]. Pixel-wise fitting of a mathematical decay model to the bleaching kinetics allows for image segmentation and for detecting heterogeneous bleaching of probes in subcellular organelles [10, 11]. From such a model, other parameters, such as the integrated probe intensity can be inferred, and image background can be detected and corrected for [3, 10, 12, 13]. The success of model-based bleaching analysis for the above-described applications depends on accurate modeling of the bleaching process. As the underlying photophysics can be complex and is not directly accessible from first principles, particularly not for the heterogeneous intracellular environment, the mathematical decay models used to describe photobleaching must be considered as empirical fitting functions with some mechanistic underpinning [10, 14, 15]. To account for complex bleaching mechanisms, distributions of rate constants are often invoked in the modeling process, whose mechanistic interpretation is difficult [10, 11, 16, 17]. On the other hand, multi-exponential fitting suffers from the well-known non-orthogonality of real exponential functions, making that fluorescence decays cannot be uniquely represented by the sum of exponential functions with real exponents [18]. Moreover, the fit quality of any model depends on minimizing movement of the specimen, since displacement of for example organelles during imaging will reduce fitting accuracy. This can only be partly alleviated by temporal and/or spatial filtering techniques, which comes to the price of eventual loss of resolution. A recent study combined the analysis of photobleaching characteristics with spectral unmixing and employed a non-negative matrix factorization to separate several fluorescent probes with nearly identical emission spectra [19]. This study demonstrated the potential of including fluorophore bleaching characteristics into spectral separation methods for live-cell imaging. But it was limited by the fact, that for each fluorescent probe, only a single bleaching fingerprint was considered. Many studies have shown, however, that photobleaching of fluorescent probes in complex environments like cells is heterogeneous [3, 11, 20-23]. In addition, uneven illumination and variations of refractive index can contribute to locally varying photobleaching kinetics in wide field and confocal microscopy [10, 12, 24, 25]. Together, this demands a full spatiotemporal description of the photobleaching characteristics for proper image segmentation and separation of spectrally indistinguishable fluorophores.

Dynamic mode decomposition (DMD) is a novel computational approach to extract dynamic information from large spatiotemporal datasets, such as images. Originating in fluid mechanics as a method to determine coherent flow patterns, DMD is increasingly applied in computer vision and biomedical imaging [26, 27]. For example, DMD has been used to detect video shots or to separate foreground from background in image sequences [28, 29]. It has also been used to segment images of kidneys and detect functional brain states by magnetic resonance imaging [30, 31]. Based on a spectral decomposition of the transfer or Koopman operator, DMD allows not only for detecting characteristic dynamic patterns in high-dimensional data sets, but also to dissect experimentally determined dynamics into individual dynamic modes. In this study, it is shown, how DMD can be employed to determine photobleaching characteristics of fluorescent probes and to distinguish probe fluorescence from cellular autofluorescence. This paper is organized as follows; first, the theory behind DMD is briefly reviewed, second, the method is applied to synthetic data of differently bleaching regions in simulated images. Third, an example of bleaching analysis in intact animals is given, where it is shown, that the characteristic photobleaching of DHE can be detected by DMD and distinguished from cellular autofluorescence in C. elegans. Fourth, it is demonstrated that DMD of the distinct photobleaching kinetics of two green-emitting fluorescent probes can be used to distinguish them in living cells. Specifically, an NBD-tagged sphingolipid and Alexa488-tagged transferrin (Alexa488-Tf), an endocytosis marker, are employed and DMD of their photobleaching kinetics allows for their unequivocal assignment to different subcellular organelles. Finally, the findings are discussed and brought into perspective for future applications of DMD in image analysis for life science applications.

## Results

### Theory of DMD applied to fluorescence photobleaching

Let *I*(*x, y, t*) be a video image sequence of fluorescence intensity, I, as a function of spatial coordinates, *x, y* and time, *t*. For the purpose of this study, this image sequence is supposed to contain the spatial distribution of some fluorescence probe in a living specimen, in which the fluorescence intensity decays over time due to photobleaching. Images have been acquired at discrete and constant time intervals, Δt, given by the microscope acquisition time. One can express each image of dimension *u times v* of the entire video sequence as *u∙v∙*1 column vectors, 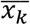, with *n* = *u∙v* entrances each, which represent the individual pixel values. Those column vectors can then be arranged in an *n × m* data matrix X consisting of *m* columns, each representing one time point of the original image sequence [26, 30]:

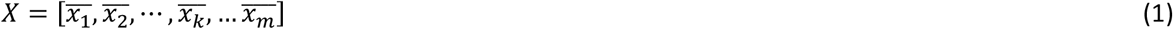

Here, the index *k*=1,…, *m* indicates the frame number resembling the time axis of the video sequence. We want to find a mapping between discrete time steps Δt from state x(k∙Δt)=x_k_ to x_k+1_ as:

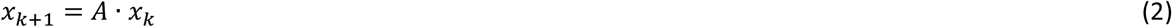

Here, A is a matrix which describes the advancement of the system in time and which resembles the Koopman or transfer operator for measurements g(*x*_k_)=*x*_k_ [27]. We want to approximate *A* solely from the given data. For that, we define the discrete time-shifted states of our system as two new matrices:

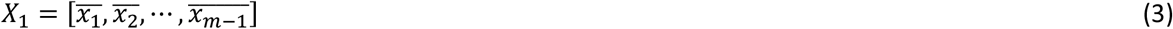

and

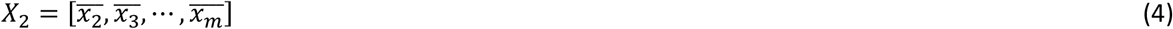

Thus, the system corresponding to Eq. (2) becomes *X*_2_ = *A*∙*X_1_*, from which we can find A by minimizing the Frobenius norm, ‖∙‖_*F*_ [27]:

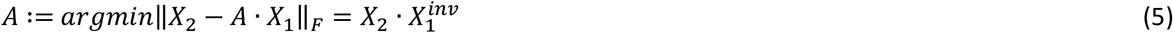

We can find the pseudoinverse of the first data matrix, 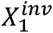, by using a singular value decomposition (SVD) of *X*_1_ into unitary matrices U, V* with singular values in the diagonal matrix Σ:

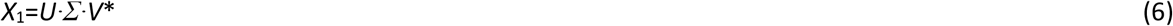

Since the data matrices, *X*_1_ and *X*_2_ have typically many more rows *n* (i.e., pixels for each image) than columns *m* (i.e., time points), there are at most *m* non-zero singular values and corresponding singular vectors, and therefore the matrix *A* will have at most rank *m*, but in practice it is calculated up to rank *r* < *m*. Thus, one can approximate *A* by calculating its projection onto these singular vectors, which gives a much smaller matrix A’ of maximal size m x m. We get [27, 28]:

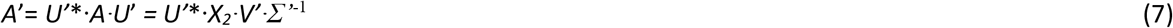

Here, *U*’, *V*’ and *Σ*’ are rank *r* ≤ *m* approximations of the full matrices, *U*, *V* and *Σ*, and * indicates the complex conjugate transpose of a given matrix (which is the transpose for a real matrix). With that, one can solve the following eigenvalue problem to determine the eigenvalues describing the temporal evolution of the studied system by spectral decomposition of A’. First, one gets the eigenvalues from the matrix Λ:

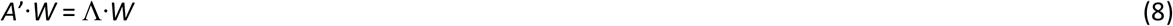

Here, the matrix W contains the corresponding eigenvectors of *A*’, providing a coordinate transformation which diagonalizes *A*’ thereby decoupling the system dynamics. Importantly, these eigenvalues and the corresponding eigenfunctions collected in the matrix, *Φ*, are the same as for the full matrix *A* [27]:

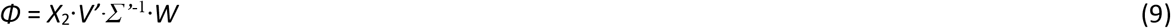

Using this spectral decomposition of the transfer matrices A and A’, respectively, we can express the dynamics of our system as linear combination of eigenfunctions, φ_j_, (i.e., eigenvectors of *A*, also called DMD modes), corresponding eigenvalues, λ_j_ (so-called DMD eigenvalues) and mode amplitudes b, which are just the components of each eigenfunction in a given direction. This leads for discrete-time systems to:

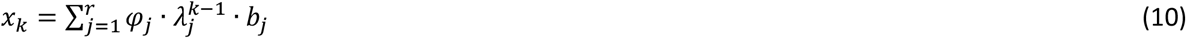

And by using the continuous eigenvalues *ω* = log(*λ*/Δ*t*), Eq. (10) can be written as:

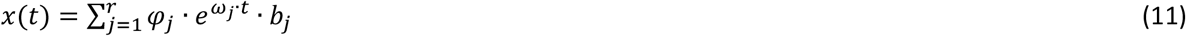

Here, x(t) is a vector of images (x, y index omitted for brevity) as a function of time, t, describing the time evolution of each dynamic mode, *φ*_j_, which is a function of time, only. The mode weights, *b*_j_, are spatial weighting matrices, i.e., functions of pixel coordinates (x, y). By this approach, the spatiotemporal image data can be decomposed in a purely data-driven manner, revealing dynamic properties of the system. In the following it is shown, how these dynamic modes can be calculated for image stacks containing bleaching fluorophores and autofluorescence respectively.

### DMD of synthetic bleach stacks

To assess the potential of DMD to capture the dynamics of photobleaching in microscopy images, synthetic image stacks with known bleach rates were used. Bleach rate constants were set to *k* = 0.1 s^-1^, 0.05 s^-1^ and 0.01 s^-1^, respectively (Fig. 1). Thus, the three circles cannot be distinguished by their intensity, but only by their different bleaching kinetics. We showed previously, that pixel-wise bleach rate fitting is able to segment these circles and recover precise estimates of the bleach rate constants for varying noise levels [8].

**Figure 1.**
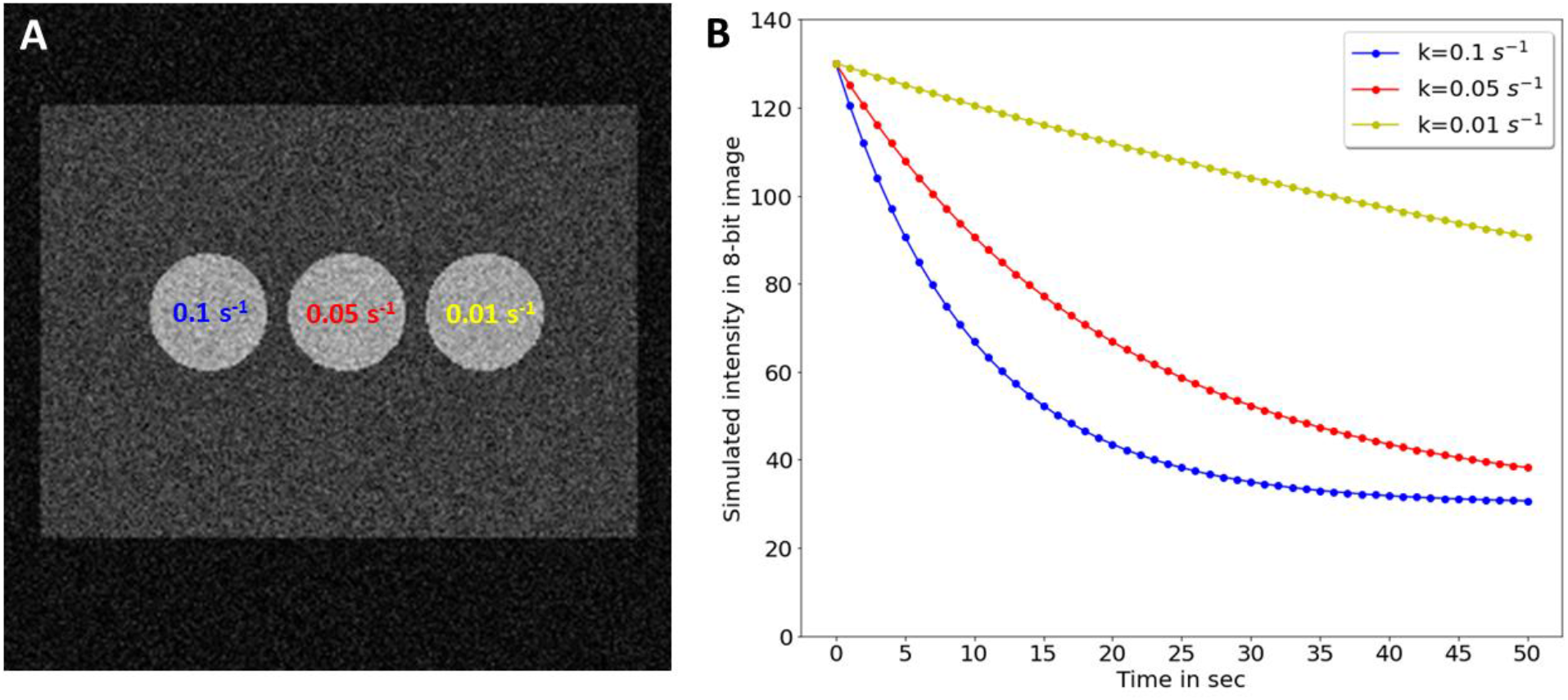
Synthetic images with regions of identical intensity but differing bleaching kinetics. A synthetic image stack was generated in which three circular regions with mean intensity = 100 reside on a rectangular area of mean intensity = 30 and bleach with the indicated rate constants of *k* = 0.1 s^-1^ (blue, left circle), *k* = 0.05 s^-1^ (red, middle circle) and *k* = 0.01 s^-1^ (yellow, right circle). A, first frame of the image stack; B, decaying mean intensity of each circle with decay rate constants color-coded as in panel A. The frame rate was 1 Hz.

To dissect the bleaching dynamics in the image stacks, DMD of rank 3 was employed. While dynamic mode 1 (‘Mode 1’) has a negative amplitude and negative mode weights, dynamic modes 2 and 3 decay exponentially with positive amplitude and weights (‘Mode 2’ and ‘Mode 3’, Fig. 2). The imaginary part of all three mode weights is zero, since there are no oscillations in the signal and no lateral displacement of the sample. There are three real eigenvalues, one for each mode (Fig. 2E). One of those is almost equal to one (i.e., λ_1_ = 0.9987), corresponding to ω_1_ = − 0.00133 upon logarithmic scaling (see Eq. 12), indicating that the corresponding dynamic mode decays very slowly. The second eigenvalue is slightly smaller, (i.e., λ_2_ = 0.9218, ω_2_ = − 0.08145) and the third is the smallest (i.e., λ_2_ = 0.856, ω_2_ = − 0.1554), all describing the time-decaying bleaching processes. Note, that these eigenvalues are not expected to resemble the rate constants used in the simulations, since DMD is not a model fitting technique. Instead, the dynamics at each pixel position is described as the sum of all three dynamic modes multiplied by the exponentials and mode weights (see Eq. 11). Importantly, the dynamics of this simulated bleach stack can be reconstructed very well with the additional benefit, that most noise is removed, such that the three circles can be better discerned than in the simulated original data (Fig. 3). Thus, DMD can be used to denoise the bleach stacks, which improves the ability to segment regions based on bleaching kinetics.

**Figure 2.**
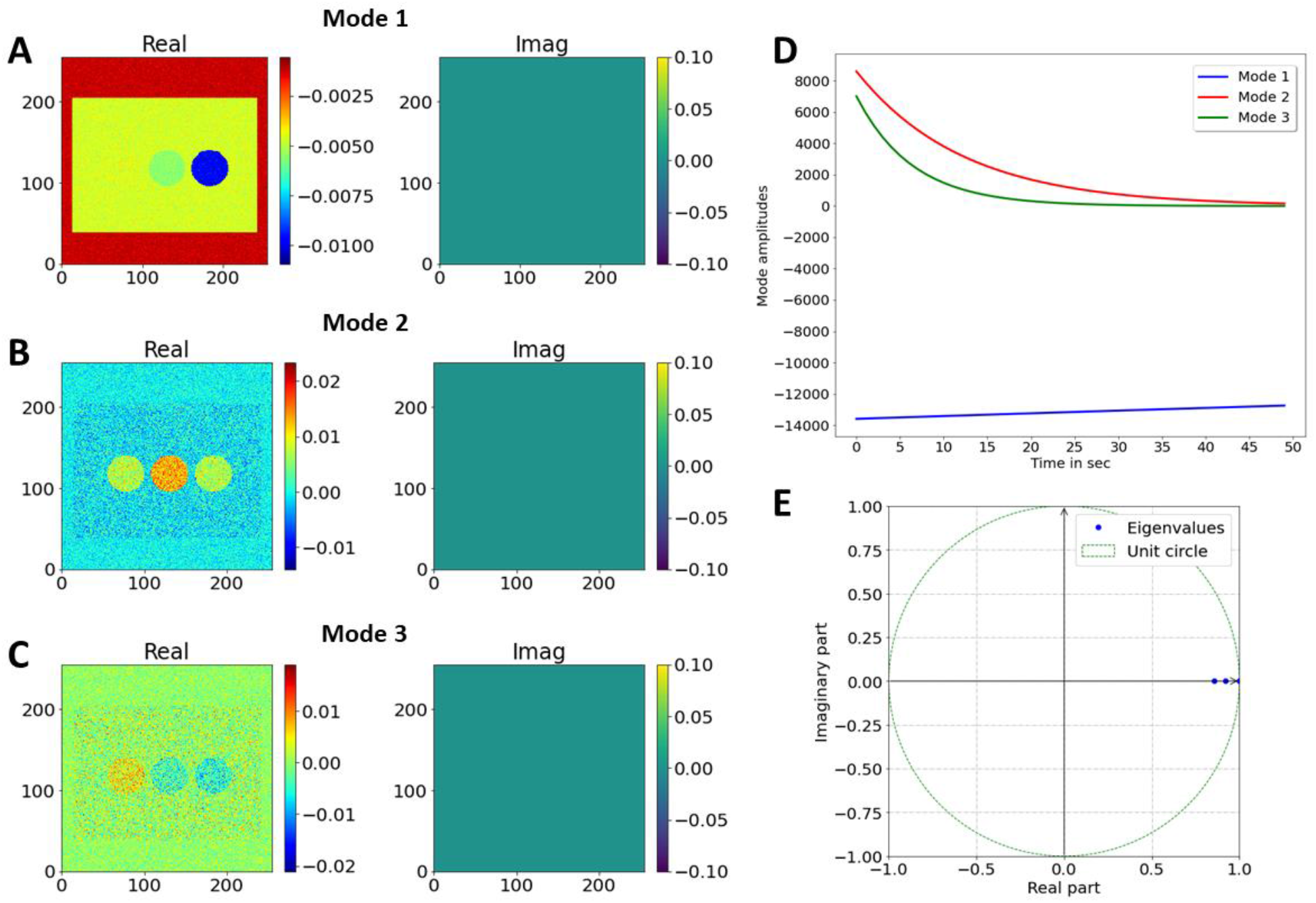
Dynamic mode decomposition of synthetic bleach stacks. A rank-3 approximation of the full matrix A was employed to decompose the simulated bleaching kinetics. A-C, mode weights for DMD mode 1 (A), DMD mode 2 (B) and DMD mode 3 (C). Real part of mode weights is shown in left panels (‘Real’), while imaginary parts are shown in right panels (‘Imag’). Only the real parts have non-zero entries. D, mode amplitudes plotted as function of time; E, eigenvalues ω1 to ω3 plotted on the unit circle.

**Figure 3.**
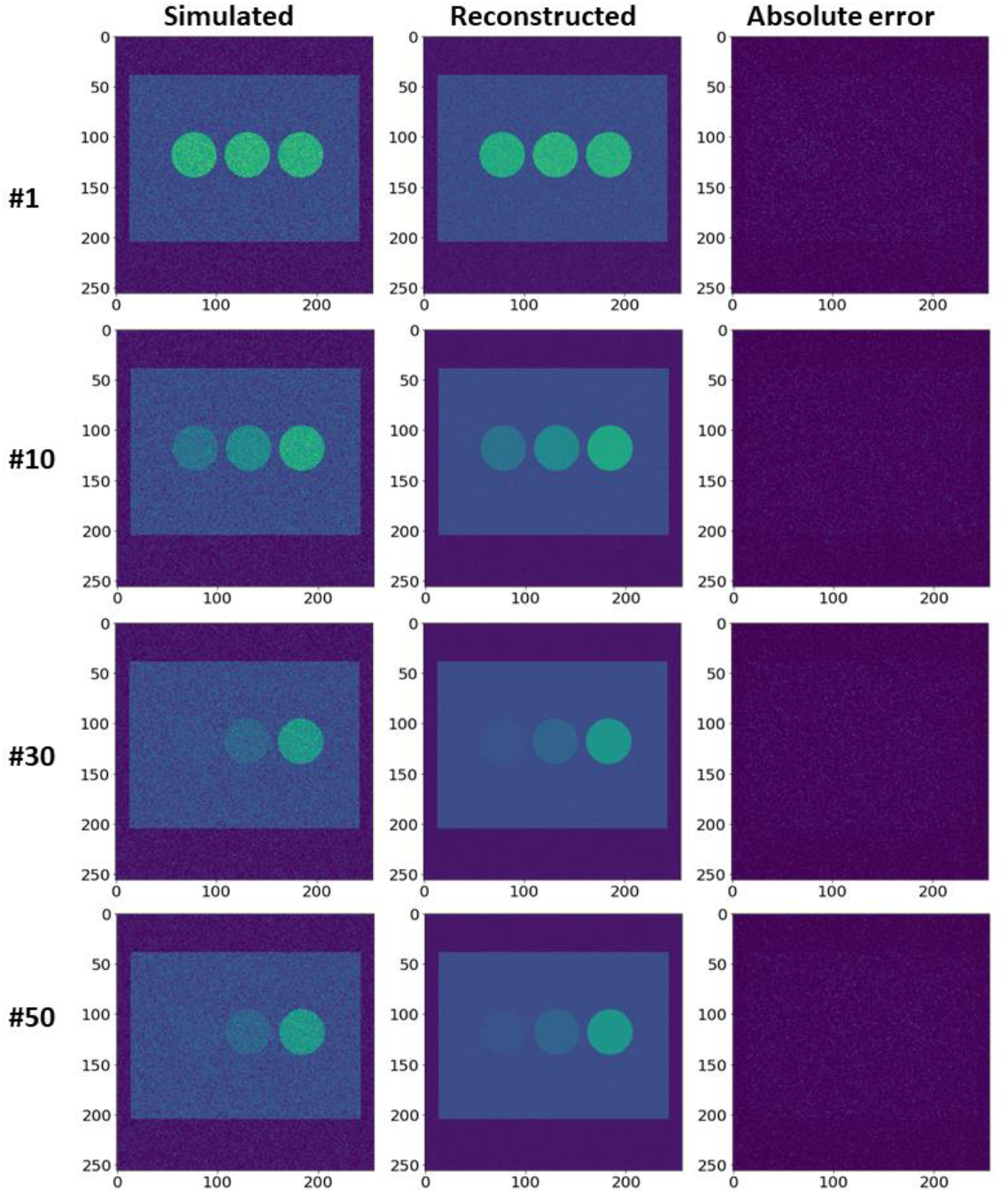
Comparison of simulated and reconstructed bleach stacks. Selected frames (#1, #10, #30 and #50) are plotted for the simulated bleach stack (left column) and the reconstructed image stack obtained from the DMD of the synthetic image series (middle column). Right column, absolute error between simulation and reconstruction. The intensity range is color-coded between 0 and 255 intensity units. One can see, that DMD approximates the simulated images very well and is also efficient in removing image noise.

### DMD of experimental image stacks of photobleaching in C.elegans

To assess, whether DMD can be used to segment real experimental data, C. elegans nematodes labeled with the intrinsically fluorescent sterol DHE were repeatedly imaged. Our earlier studies showed that DHE’s fluorescence overlaps strongly with autofluorescence of nematodes in the ultraviolet region of the spectrum, but also that DHE bleaches much faster than autofluorescence [8, 9]. To decompose the differential bleaching of probe and autofluorescence, image stacks were analyzed by DMD using a rank-5 approximation of the full data matrix. Strikingly, the reconstructed image stack containing the information from all five dynamic modes perfectly matches the original data, and the decreasing integrated intensity of both image stacks coincides closely (Fig. 4).

**Figure 4.**
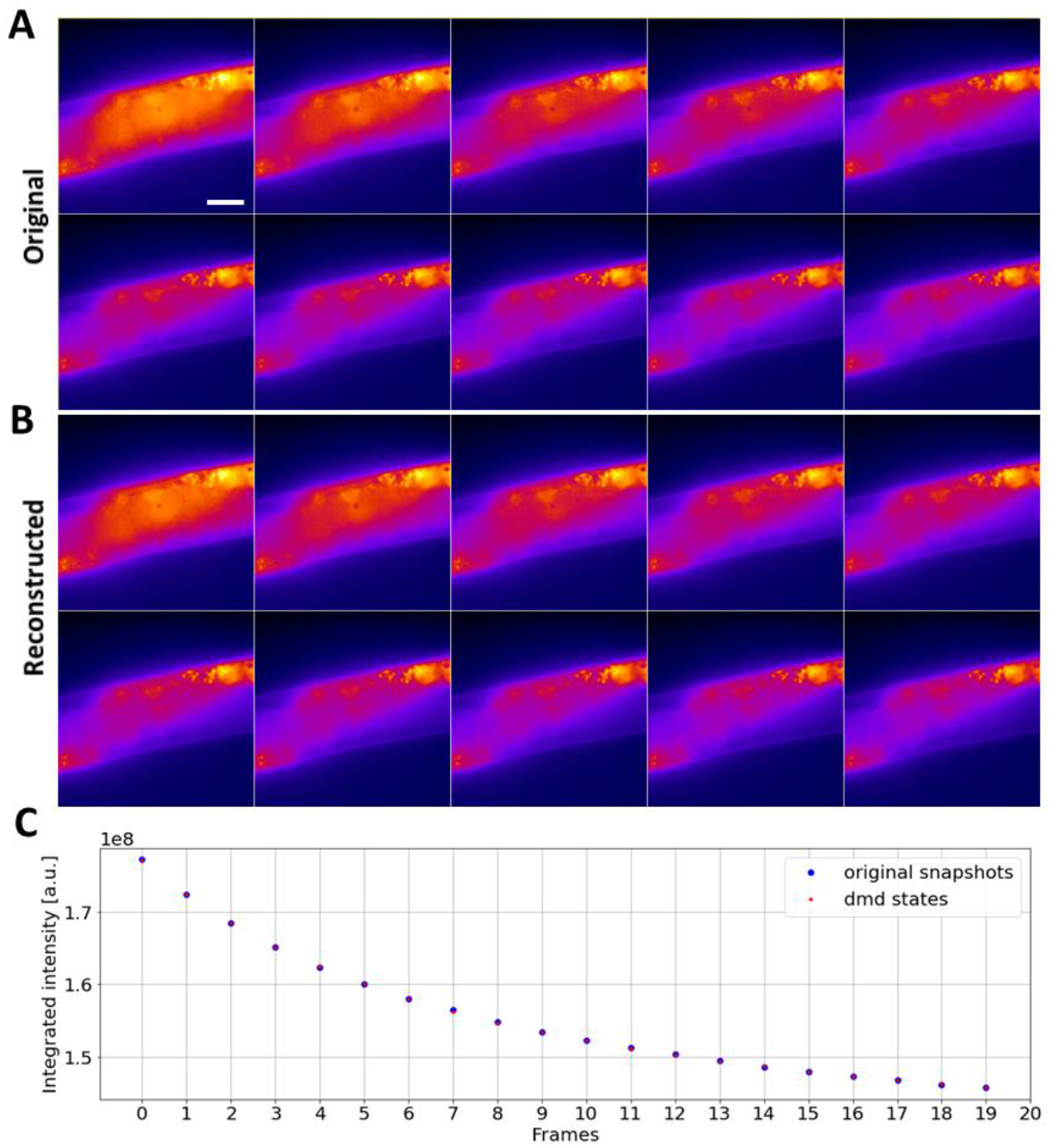
Comparison of experimental and reconstructed bleach stacks of DHE in C. elegans. A, B montage of selected frames (every 4^th^ frame) of the experimental fluorescence stacks of DHE labeled C. elegans (A) and of the reconstructed image stack obtained from the DMD of rank 5 of the experimental image series (B). A FIRE LUT is chosen to color-code intensities. C, integrated intensity (i.e., sum of pixel values per frame) for the experimental data (blue circles) and the DMD reconstruction (red circles). Bar, 20 μm.

DMD of this fluorescence image series resulted in two complex eigenvalues, which describe two degenerated modes with oscillating dynamics (Fig. 5A, ‘Mode 1 and 2’, and C). In addition, there are three real eigenvalues smaller than one, corresponding to decaying dynamic modes, which describe the bleaching kinetics of DHE and autofluorescence respectively (Fig. 5B ‘Mode 3 to 5’, and C).

**Figure 5.**
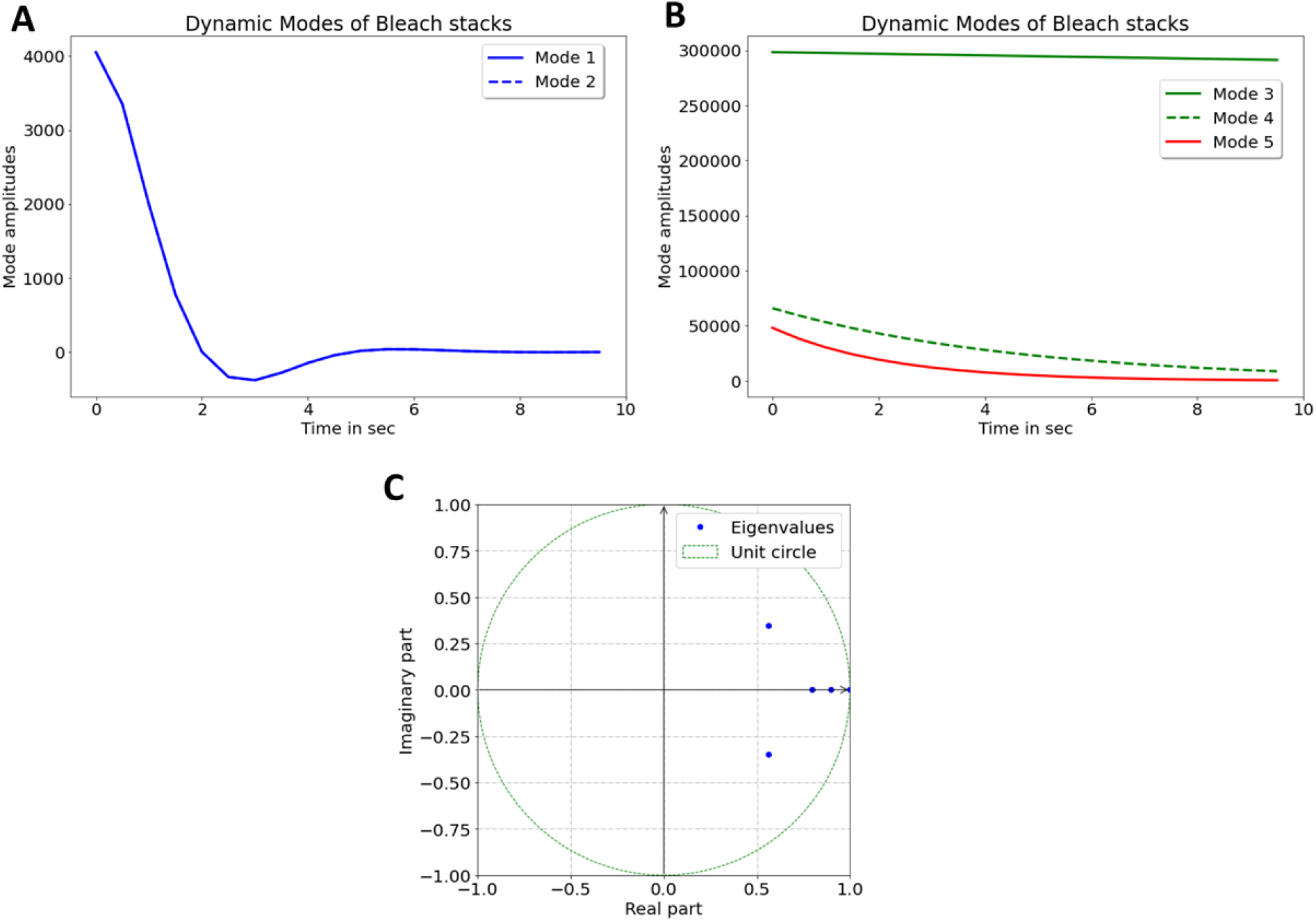
Dynamic mode amplitudes and eigenvalues of fluorescence images of DHE-labeled nematodes. A, B, mode amplitudes of a rank-5 DMD of the experimental bleach stacks of DHE labeled C. elegans with two oscillatory modes (Mode 1 and 2, A) and three exponentially decaying amplitudes (Mode 3 to 5, B). C, eigenvalues ω_1_ to ω_5_ plotted on the unit circle. The first two eigenvalues have non-zero imaginary part and are degenerate (see also the corresponding oscillatory amplitudes in A). Eigenvalues 3 to 5 are real and smaller than one, describing decaying intensities in the bleach stacks.

Inspection of the mode weights confirms this interpretation (Fig. 6); Mode 1 and 2 contain non-zero imaginary parts, which account for the slight lateral displacement of the nematodes during imaging (Fig. 6A and B). Such movement cannot be entirely prevented despite anesthetizing the animals before imaging, and it can impact the fit quality in pixel-wise bleach rate fitting [8]. In contrast, in DMD one can account for some movement of subcellular structures without compromising the bleaching analysis. Mode 3 to 5 describe the decaying intensity, as inferred from the mode weights which all have only real entries (Fig. 6C-E). These modes have real eigenvalues, smaller than one and decaying mode amplitudes (Fig. 5B and C).

**Figure 6.**
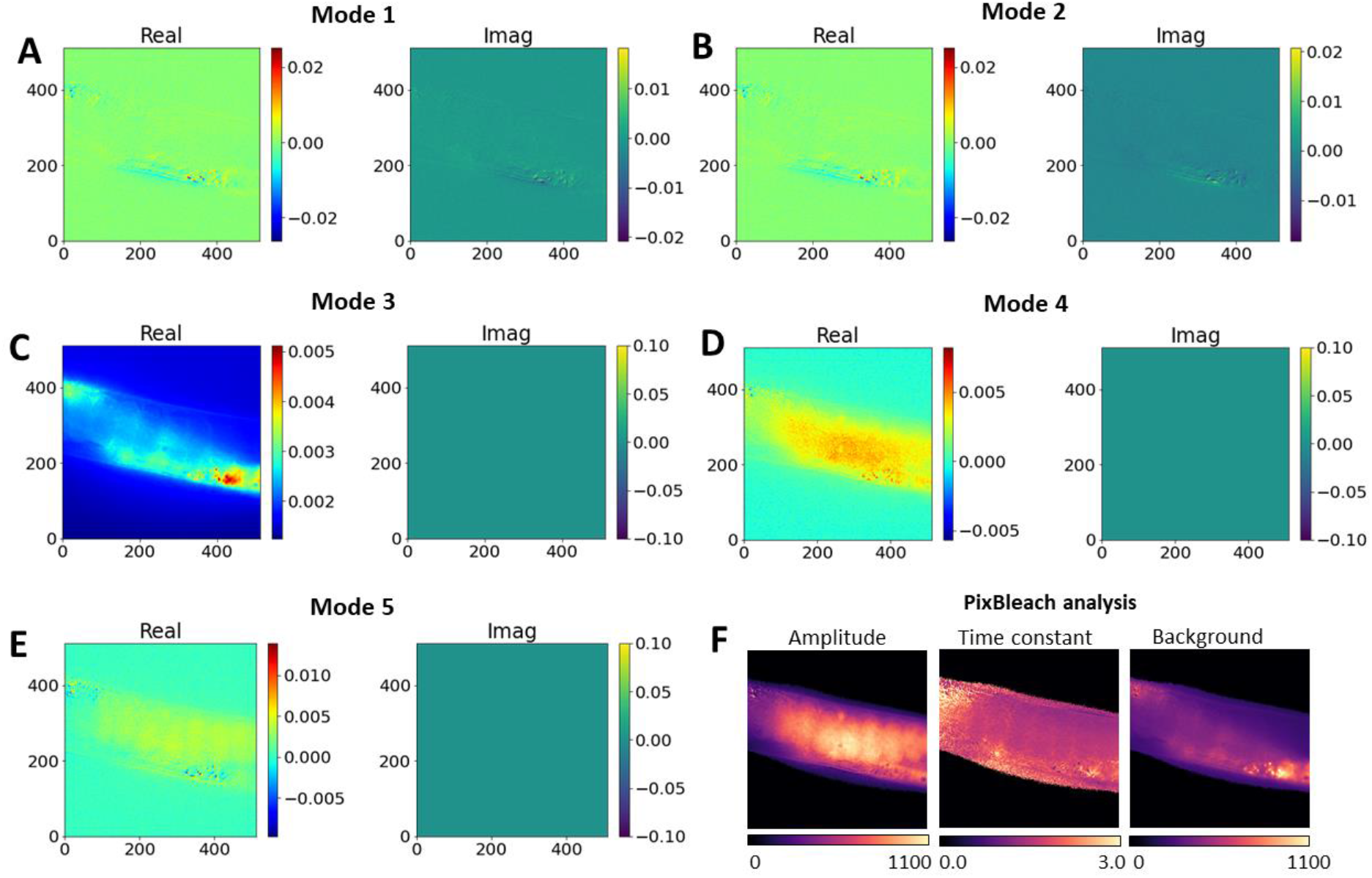
Mode weights of DMD of C.elegans bleach stacks and comparison with pixel-wise fitting. A-E, mode weights for DMD modes 1 to 5. The real part of mode weights is shown in left panels (‘Real’), while the imaginary parts are shown in right panels (‘Imag’). F, bleach rate fitting using a stretched exponential function with bleaching amplitudes (right panel), time constant map (middle panel) and background fluorescence (right panel). The amplitude image in F shows the distribution of the rapidly bleaching DHE, while the background image resembles most of the autofluorescence.

Mode 3 described the slowly decaying autofluorescence, while Mode 4 and 5 describe bleaching of the fluorescent sterol DHE. This interpretation is supported by comparing the outcome of DMD with bleach-rate fitting, which shows that significant bleaching takes place in the region of the oocytes and intestinal cells, where the DHE resides (see ‘Amplitude’ image in Fig. 6F), closely resembling the mode weight images for Mode 4 and 5 (Fig. 6D and E). We and others showed previously that oocytes are particularly sterol-rich cells in C. elegans, since the ingested sterols are essential for steroid hormone production to control developmental transitions [7, 8, 32, 33]. Pixel-wise bleach rate fitting identifies autofluorescence as non-bleaching, i.e., constant background term (Fig. 6F). That background map is very similar to the real peart of the weight image of Mode 3 of the DMD (compare Fig. 6C, left panel and F, right panel).

Having decomposed the entire dynamics in the bleach stacks of nematodes, DMD allows for separate inspection and analysis of each dynamic mode. This makes it possible to separate autofluorescence from DHE intensity and thereby to segment the images into different fluorescence contributions. By using the logarithmically weighted eigenvalues, the mode weights, and the mode amplitudes, one can calculate image stacks representing the individual dynamic modes, representing a decomposition of the entire bleaching dynamics in the original image stacks (see Eq. 11 and 12).

From that, one can clearly infer cellular autofluorescence from Mode 3 and more rapidly decaying DHE fluorescence from Mode 4 and 5, respectively (Fig. 7). Clearly, the sum of Mode 4 and 5 resembles the total DHE fluorescence, which bleaches much faster than cellular autofluorescence. Accordingly, DMD allows for segmentation of image structures based on their dynamics, which makes it possible that one pixel contains information from both dynamic structures, just to different extend, as visualized in a color overlay of the decomposed bleaching dynamics (Fig. 7B-D). This is not straightforward to implement in pixel-based fitting, where pixel overlap of two regions can only be accounted for by bi-exponential fitting, which often fails, when the signal-to-noise ratio in image regions gets to small [8]. Here, DMD is more robust and allows for accurate segmentation since there are no model parameters to be inferred from the data. In addition, DMD but not pixel-wise fitting provides the bleaching dynamics of each fluorescent entity as separate image stacks.

**Figure 7.**
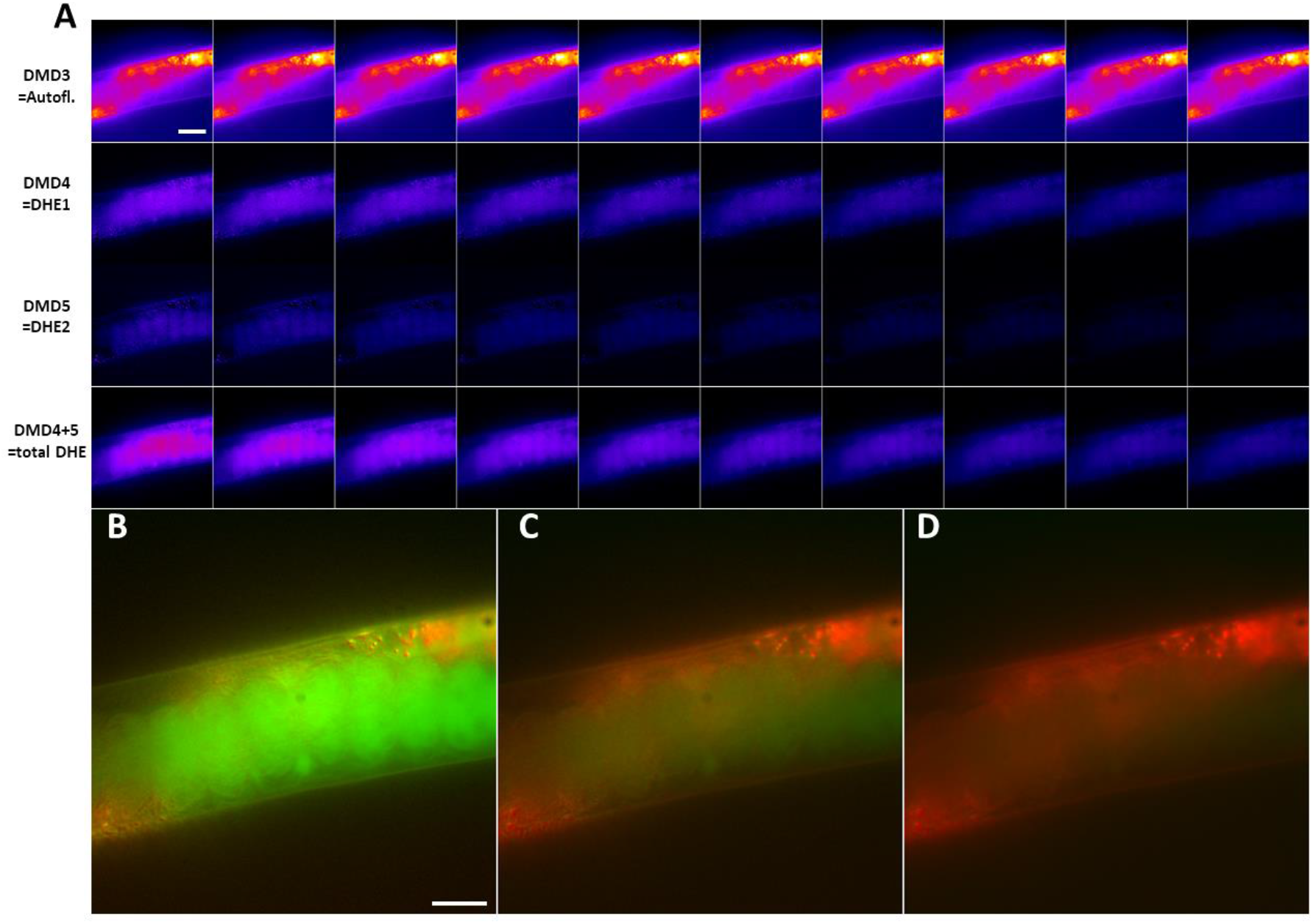
DMD of C. elegans bleach stacks allows for discrimination of autofluorescence from DHE probe intensity. A, montage of individual dynamic modes shown as every second image of the corresponding image stacks. Dynamic mode 3 (‘DMD3’) resembles cellular autofluorescence of nematodes (upper row in A), while dynamic mode 4 and 5 (‘DMD4’ and ‘DMD5’) constitute DHE fluorescence (two middle rows in A). The sum of mode 4 and 5 shows the total DHE fluorescence (lower row in A). B-D, color overlay of mode decomposition with dynamic mode 3 resembling autofluorescence in red and sum of mode 4 and 5 representing DHE fluorescence in green. B, first frame, C, 10^th^ frame and D, 20^th^ frame of this color representation of the DMD, showing the rapid bleaching of DHE fluorescence (green) compared to autofluorescence (red). Bar, 20 μm.

### DMD of experimental image stacks of mammalian cells labeled with two green-fluorescent probes

To further explore the potential of the method, DMD was used to discriminate two fluorescence tagged molecules, which are spectrally indistinguishable, but show different bleaching kinetics in living cells. BHK cells were labeled with two green emitting fluorescence probes; the iron-transporting protein transferrin tagged with an Alexa488-dye, which is very photostable (Fig. S1), and the membrane probe C6-NBD-SM, which bleaches much faster (Fig. S2). Alexa488-Tf will bind to the transferrin receptor and become internalized by clathrin-mediated endocytosis followed by recycling from early sorting and recycling endosomes [34]. The latter is also called the endocytic recycling compartment (ERC) and appears as perinuclear enrichment of small vesicles in Chinese hamster ovarian (CHO) and BHK cells [35, 36]. C6-NBD-SM is a fluorescent sphingolipid probe, which has been shown to be targeted to the ERC and recycled from the cell with very similar kinetics as Tf, which is why this fluorescent lipid probe is often seen as ‘bulk membrane recycling marker’ [35, 37-39]. Thus, both probes accumulate in the perinuclear ERC, but to different extent. While the majority of C6-NBD-SM remains in the plasma membrane (PM), almost the entire pool of Alexa488-Tf will accumulate in the ERC in this experiment. By loading C6-NBD-SM onto albumin, the lipid probe can be rapidly inserted into the PM, followed by its endocytosis and trafficking through the endocytic recycling pathway together with fluorescent Tf [39]. Such co-trafficking is normally assessed in two-color fluorescence imaging experiments, in which the emission color of the fluorescent probes is separated using suitable filter combinations [37-39]. Using different colors for two endocytic markers limits the number of additional channels to be available for other probes to two on most wide field and confocal microscopy systems (e.g., there is typically an additional blue filter set for DAPI and an infrared filter cube for another organelle marker).

By decomposing the different bleaching kinetics of the two green emitting endocytic probes, Alexa488-Tf and C6-NBD-SM using DMD, their intracellular distribution can be determined using only one filter set (Fig. 8). As shown in Fig. 8A, the green fluorescence bleaches rapidly in the PM, where a major portion of C6-NBD-SM resides but much slower in the ERC, where the majority of the more photostable Alexa488-Tf is located. This notion is sustained by DMD of image stacks of cells labeled with only Alexa488-Tf (Fig. S1) or only with C6-NBD-SM (Fig. S2). The heterogeneous bleaching dynamics of doubly labeled cells can be decomposed by DMD into five dynamic modes, three of which show fast decay (Mode 1, 3 and 4; Fig. 8C). Mode 2 is almost constant and has an eigenvalue close to one (, Fig. 8D), while Mode 5 decays slowly (dashed blue and straight red line in Fig. 8C). From the corresponding weight amplitudes, one sees that Mode 1, 3 and 4 have intensity in the PM and in the ERC, while Mode 2 and 5 have non-zero intensities almost exclusively in the perinuclear area (Fig. S3). Based on these observations, the sum of Mode 1, 3 and 4 are assigned to the lipid marker C6-NBD-SM, while the sum of Mode 2 and 5 are assigned to Alexa488-Tf. The intracellular distribution of C6-NBD-SM and Alexa488-Tf overlaps in the perinuclear ERC but only very little the PM (see color overlay in Fig. 8B with C6-NBD-SM in green and Alexa488-Tf pseudo-colored in red). Clearly, while C6-NBD-SM bleaches rapidly (Fig. 8B, green), Alexa488-Tf does bleach as well but much more slowly (Fig. 8B, red). Both conclusions are confirmed from stacks of single-labeled cells (Fig. S2 and S3). In pixel-wise bleach rate fitting one cannot separate the rapidly bleach lipid probe from the slowly bleaching Alexa488-Tf, which instead would be assigned to the background term (Fig. S3F). These results demonstrate the potential of DMD in decomposing photobleaching dynamics for efficient separation of different fluorophores in live-cell microscopy.

**Figure 8.**
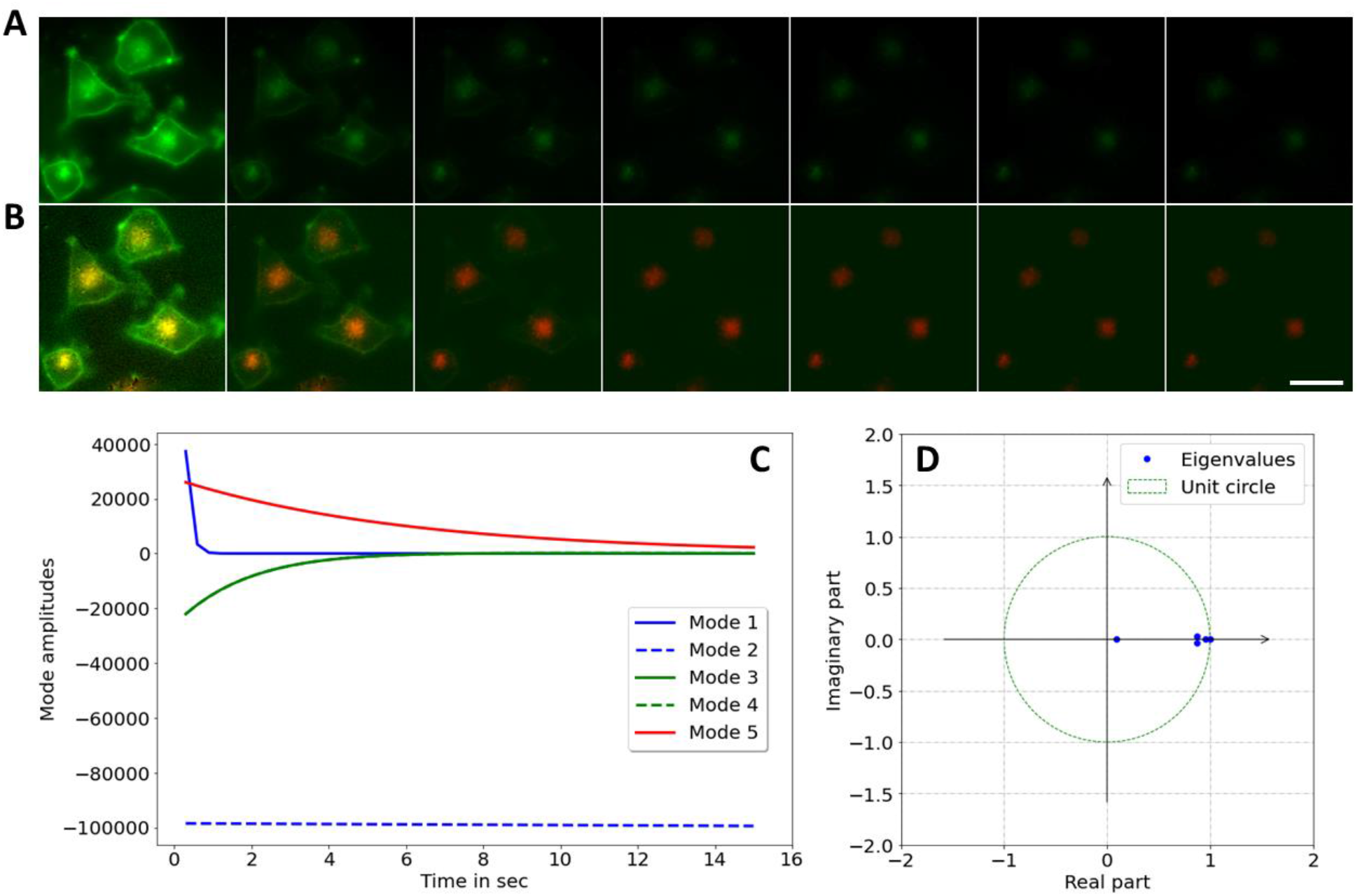
DMD of bleach stacks of BHK cells labeled with two green probes, Alexa488-Tf and C6-NBD-SM. BHK cells were labeled with 4 μM C6-NBD-SM and with 20 μg/ml Alexa488-Tf, both emitting in green, as described in Materials and Methods. A, montage of selected frames (every 5th frame) of such double labeled cells. B, reconstruction of DMD of the bleach stack in A with the sum of dynamic modes 1, 3 and 4 shown in green (resembling the fast-bleaching C6-NBD-SM) and sum of dynamic mode 2 and 5 (resembling the slowly bleaching Alexa488-Tf) shown in red. Bar, 20 μm. C, mode amplitudes and D, eigenvalues of the DMD plotted on the unit circle.

## Discussion

In this study, it is shown that image segmentation and separation of spectrally indistinguishable fluorophores in live-cell microscopy can be achieved by DMD of their photobleaching kinetics. This is demonstrated first on synthetic image stacks with simulated photobleaching and thereafter on two different experimental data sets. In all cases, DMD can decompose the photobleaching kinetics properly, allowing for clear separation of probe from autofluorescence or of different fluorescent dyes in the same sample. One can therefore envision, that DMD of probe photobleaching can be combined with spectral unmixing, thereby increasing the number of probes which can be separately detected in parallel [1, 19]. Additionally, DMD can compensate for noise and small movement artefacts, making it a powerful computational tool for analyzing photobleaching in live-cell imaging experiments in the future. There is a strong interest in developing novel fluorescent probes for biomedical imaging applications. Here, a concern is often that prolonged fluorescence imaging of dynamic processes in living cells can lead to photo destruction of fluorophores [40]. Therefore, it is important to determine photobleaching characteristics of fluorescent dyes under various conditions, either for optimizing probe design or for optimizing imaging conditions [14, 20, 41, 42]. For example, sensitive single molecule imaging and super-resolution microscopy critically depends on development of bright and photostable fluorophores, and here, DMD of dye photobleaching in cells can be very useful for designing improved organic fluorophores [43]. DMD will also be useful in discriminating different fluorophores based on their characteristic bleaching propensities, for multi-color super-resolution microscopy [44]. Similarly, optimizing photosynthetic antenna systems for light harvesting applications in photovoltaics demands photostable pigment structures, and analysis of their photo destruction using imaging-based DMD can be combined with other approaches such as lifetime and absorption measurements as well as electron paramagnetic resonance spectroscopy in the future [45].

Furthermore, photobleaching kinetics also report about environmental impacts on a fluorophore, since in many cases, photobleaching kinetics is inversely related to the fluorescence lifetime of a fluorophore. For example, if a nearby acceptor molecule can receive the energy of an excited fluorophore due to Förster resonance energy transfer (FRET) its excited lifetime gets reduced and accordingly its bleaching propensity lowered, since photobleaching can only take place from an excited state. Therefore, photobleaching kinetics of the donor molecule can report about FRET efficiencies, similar as lifetime imaging [22, 46, 47]. Under conditions, where lifetime imaging is not feasible or at least very difficult, e.g., for weakly emitting UV probes, analysis of donor photobleaching kinetics is a very useful approach, and here DMD is a good tool for their analysis. The same applies to quenching studies, in which a dynamic quencher is used to determine the accessibility of a fluorescent probe, for example when analyzing the permeability of the fungal cell wall [48], or the transbilayer distribution of a membrane probe between the two PM leaflets [49-52]. Dynamic quenching shortens the fluorescence lifetime and thereby slows the photobleaching of the quenched dye [10, 13, 53], which can be detected by DMD of its photobleaching kinetics. The extent of photobleaching is often directly proportional to the occupation of the triplet state of a fluorophore, from which reaction with singlet oxygen and thereby photooxidation can take place. This concept is used in photodynamic therapy, where light-induced production of reactive oxygen species is used to kill tumor cells selectively [54-56]. Here, control of the photobleaching process is essential, and DMD of photobleaching kinetics can become a useful tool in its analysis.

## Conclusions

A new computational method is presented to analyze photobleaching kinetics of fluorescent entities in microscopy images of living cells in a purely data-driven manner. It is shown that the decomposition of photobleaching kinetics into dynamic modes allows for image segmentation, image denoising and discrimination of different fluorescent probes and autofluorescence on a pixel-by-pixel basis. This novel approach can be combined with spectral unmixing and FRET studies, for example as part of large-scale image-based screens to assess organelle and marker distribution in multi-color experiments.

## Declarations

### a. Ethics approval and consent to participate

Not applicable.

### b. Consent for publication

Not applicable.

### c. Availability of data and materials

All data analyzed during this study are included in this published article [and its supplementary information files]. The datasets generated and/or analyzed during the current study are also available in the GITHUB repository, (https://github.com/DanielW-alt/Photobleaching).

### d. Competing interests

I declare that the author has no competing interests as defined by BMC, or other interests that might be perceived to influence the results and/or discussion reported in this paper.

### e. Funding

DW acknowledges funding from the Villum foundation (grant nr. 73288).

### f. Authors’ contributions

DW is the sole author of this manuscript, so DW has carried out the work described in this study and has made the figures and written and reviewed the manuscript.

## g. Acknowledgments

DW acknowledges technical assistance from Tanja Christensen, BMB, SDU, Denmark.

## Materials and Methods

### Materials

N-[6-[(7-nitro-2-1,3-benzoxadia-zol-4-yl)amino]-dodecanoyl]-sphingosine-1-phosphocholine (C6-NBD-SM) was obtained from Avanti Polar Lipids (Alabaster, AL). DHE was purchased from SIGMA Chemical (St. Louis, MA). Buffer medium contained 150mM NaCl, 5mM KCl, 1mM CaCl2, 1mM MgCl2, 5mMglucose, and 20mMHEPES (pH 7.4). Alexa488-protein labeling kit was purchased from Molecular Probes (ThermoFisher). Transferrin was iron loaded as described previously [57]. The succinimidyl ester of Alexa-588 was conjugated to the iron-loaded transferrin to get Alexa488-Tf following the manufacturer’s instructions. C6-NBD-SM was loaded onto fatty-acid free bovine serum albumin (BSA) following our previously published procedure [58]. BHKasc cells were kindly provided by Dr. Kirsten Sandvig, Cancer Center, Norwegian Radiation Hospital, University of Oslo, Norway.

### Labeling and imaging of nematodes

Wild-type C. elegans strains were cultured, labeled with DHE and imaged on a UV-sensitive wide field microscope, exactly as described previously [8]. Images of nemtatodes were acquired in the UV channel (335-nm (20-nm bandpass) excitation filter, 365-nm dichromatic mirror and 405-nm (40-nm bandpass) emission filter) with an acquisition time of 500 msec and no pause between acquisitions.

### Culture, labeling and imaging of Baby hamster kidney (BHK) cells

BHKasc cells were grown in DMEM supplemented with 7.5% heat-inactivated FCS, 2mM l-glutamine, 100 units/ml penicillin, 100_g/ml streptomycin, 0.2 mg/ml geneticin, and 2 μg/ml tetracycline [59]. Three days prior to the experiments, the cells were seeded on microscope slide dishes and kept in the same medium until the experiments. Cells were labeled with 4 μM C6-NBD-SM for 5 min at 37° C and washed three times with buffer medium. Subsequently, cells were labeled with 20 μg/ml Alexa488-Tf for 30 min at 37° C and washed three times with buffer medium. In separate control experiments, cells were either labeled just with C6-NBD-SM, washed and chased for 30 min or just with Alexa488-Tf for 30 min at 37° C and washed three times with buffer medium before imaging.

### Image simulation and data analysis

To validate the procedure synthetic bleach stacks with known bleaching characteristics were generated using the Macro language of ImageJ (https://imagej.nih.gov/ij/), as described previously [8, 60]. Specifically, an 8-bit image stack with a background of random intensities with mean intensity equal to 30 was generated in which a rectangular region of mean intensity 60 contained three circles with mean intensity of 160 each. DMD and accompanying analysis was carried out in Python using Jupyter notebooks (https://jupyter.org/) and PyDMD, a python library for DMD calculations [61]. In brief, upon SVD of the image data matrix with either a pre-defined rank or matrix-specific optimal rank, the DMD modes are calculated. Eigenvalues determined for a rank-*r* decomposition of the system matrix *A*, λ_j_, for *j*=1,…, *r*, are logarithmically scaled and divided by the interval time (i.e. the acquisition time in the case of bleach stacks, Δt):

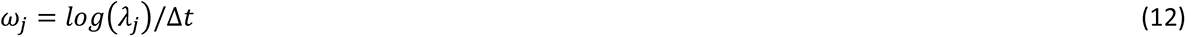

Using the calculated dynamic modes, *φ*_j_, and their weights *b*_j_ the time evolution of each dynamic mode is calculated according to Eq. 11, above. For pixel-wise bleach rate fitting a plugin to ImageJ, PixBleach, was used [8, 62]. Both methods were compared by calculating the root mean square error (RMSE) between data and model.

## Supplementary figure legends

**Additional File 1: Figure S1. Dynamic mode decomposition of image stacks containing Alexa488-Tf labeled cells.** BHK cells were labeled with 20 μg/ml Alexa488-Tf for 30 min, washed with buffer medium and imaged on a wide field fluorescence microscope. A, selected frames of an image stack acquired with 0.3 sec acquisition time and without pause. Images are identically scaled; bar 10 μm. B, C, DMD of this image stack using a rank-5 approximation to the full transfer matrix. B, mode weights and C, mode amplitudes as function of time.

**Additional File 2: Figure S2. Dynamic mode decomposition of image stacks containing C6-NBD-SM labeled cells.** BHK cells were labeled with 4 μM C6-NBD-SM for 30 min, washed with buffer medium and imaged on a wide field fluorescence microscope. A, selected frames of an image stack acquired with 0.3 sec acquisition time and without pause. Images are identically scaled; bar 10 μm. B, C, DMD of this image stack using a rank-5 approximation to the full transfer matrix. B, mode weights and C, mode amplitudes as function of time.

**Additional File 3: Figure S3. Mode weights for DMD of image stacks of BHK cells labeled with C6-NBD-SM and Alexa488-Tf.** BHK cells were labeled with 4 μM C6-NBD-SM and with 20 μg/ml Alexa488-Tf, both emitting in green, as described in Materials and Methods. BHK cells were labeled with 4 μM C6-NBD-SM for 30 min, washed with buffer medium and imaged on a wide field fluorescence microscope. Mode weights for DMD of rank 5 of this data are shown. The real part of mode weights is shown in left panels (‘Real’), while the imaginary parts are shown in right panels (‘Imag’). F, bleach rate fitting using a stretched exponential function with bleaching amplitudes (right panel), time constant (middle panel) and background term (left panel).

**Additional file 4: Simulated bleach stack.** Raw data set 1 used in the analysis shown in Figures 1 to 3. Photobleaching was simulated using single-exponential decay functions as described in Materials and Methods.

**Additional file 5: Experimental bleach stack of C. elegans labeled with DHE.** Raw data set 2 used in the analysis shown in Figure 4 to 7. C. elegans was labeled with DHE, and images were acquired with 0.5 sec acquisition time and without pause as described in Materials and Methods.

**Additional file 6: Experimental bleach stack of BHKasc cells double-labeled with C6-NBD-SM and Alexa488-Tf.** Raw data set 3 used in the analysis shown in Figure 8 and S3. BHKasc cells were labeled with C6-NBD-SM and Alexa488-TF, and images were acquired with 0.3 sec acquisition time and without pause as described in Materials and Methods.

**Additional file 7: Experimental bleach stack of BHKasc cells labeled with Alexa488-Tf.** Raw data set 4 used in the analysis shown in Figure S1. BHKasc cells were labeled with Alexa488-TF, and images were acquired with 0.3 sec acquisition time and without pause as described in Materials and Methods.

**Additional file 8: Experimental bleach stack of BHKasc cells labeled with C6-NBD-SM.** Raw data set 5 used in the analysis shown in Figure S2. BHKasc cells were labeled with C6-NBD-SM, and images were acquired with 0.3 sec acquisition time and without pause as described in Materials and Methods.

## Notes

### Competing Interest Statement

The authors have declared no competing interest.

## References

1. McRae TD, Oleksyn D, Miller J, Gao YR: Robust blind spectral unmixing for fluorescence microscopy using unsupervised learning. PloS one 2019, 14(12):e0225410.

2. Niehorster T, Loschberger A, Gregor I, Kramer B, Rahn HJ, Patting M, Koberling F, Enderlein J, Sauer M: Multi-target spectrally resolved fluorescence lifetime imaging microscopy. Nature methods 2016, 13(3):257–262.

3. Ghauharali RI, van Driel R, Brakenhoff GJ: Structure-oriented fluorescence photobleaching analysis: a method for double fluorescent labeling studies. J Microsc 1997, 185(3):375–384.

4. Entchev EV, Kurzchalia TV: Requirement of sterols in the life cycle of the nematode Caenorhabditis elegans. Semin Cell Dev Biol 2005, 16:175–182.

5. Mörck C, Olsen L, Kurth C, Persson A, Storm NJ, Svensson E, Jansson JO, Hellqvist M, Enejder A, Faergeman NJ et al: Statins inhibit protein lipidation and induce the unfolded protein response in the non-sterol producing nematode Caenorhabditis elegans. Proc Natl Acad Sci U S A 2009, 106(43):18285–18290.

6. Ashrafi K, Chang, F.Y., Watts, J.L., Fraser, A.G., Kamath, R.S., Ahringer, J., and Ruvkun, G.: Genome-wide RNAi analysis of Caenorhabditis elegans fat regulatory genes. Nature 2003, 421:268–272.

7. Matyash V, Geier C, Henske A, Mukherjee S, Hirsh D, Thiele C, Grant B, Maxfield FR, Kurzchalia TV: Distribution and transport of cholesterol in Caenorhabditis elegans. Mol Biol Cell 2001, 12:1725–1736.

8. Wüstner D, Landt Larsen A, Færgeman NJ, Brewer JR, Sage D: Selective visualization of fluorescent sterols in Caenorhabditis elegans by bleach-rate based image segmentation. Traffic 2010, 11(4):440–454.

9. Wüstner D, Sage D: Multicolor bleach-rate imaging enlightens in vivo sterol transport. Commun Integr Biol 2010, 3(4):1–4.

10. Wüstner D, Christensen T, Solanko LM, Sage D: Photobleaching kinetics and time-integrated emission of fluorescent probes in cellular membranes. Molecules 2014, 19(8):11096–11130.

11. Koppel DE, Carlson C, Smilowitz H: Analysis of heterogeneous fluorescence photobleaching by video kinetics imaging: the method of cumulants. J Microsc 1989, 155(2):199–206.

12. Ghauharali RI, Hofstraat JW, Brakenhoff GJ: Fluorescence photobleaching-based shading correction for fluorescence microscopy. J Microsc 1998, 192(2):99–113.

13. Hirschfeld T: Quantum efficiency independence of the time integrated emission from a fluorescent molecule. Applied Optics 1976, 15:3135–3139.

14. Eggeling C, Widengren J, Rigler R, Seidel CAM: Photobleaching of fluorescent dyes under conditions used for single-molecule detection: evidence of two-step photolysis. Anal Chem 1998, 70(13):2651–2659.

15. Widengren J, Mets Ü, Rigler R: Fluorescence correlation spectroscopy of triplet states in solution: a theoretical and experimental study. J Phys Chem 1995, 99:13368–13379.

16. Solie TN, Small EW, Isenberg I: Analysis of nonexponential fluorescence decay data by a method of moments. Biophysical journal 1980, 29(3):367–378.

17. Steinbach PJ, Chu K, Frauenfelder H, Johnson JB, Lamb DC, Nienhaus GU, Sauke TB, Young RD: Determination of rate distributions from kinetic experiments. Biophysical journal 1992, 61(1):235–245.

18. Istratov AA, Vyvenko OF: Exponential analysis in physical phenomena. Rev Sci Instrum 1999, 70(2):1233–1257.

19. Orth A, Ghosh RN, Wilson ER, Doughney T, Brown H, Reineck P, Thompson JG, Gibson BC: Super-multiplexed fluorescence microscopy via photostability contrast. Biomed Opt Express 2018, 9(7):2943–2954.

20. Song L, Hennink EJ, Young IT, Tanke HJ: Photobleaching kinetics of fluorescein in quantitative fluorescence microscopy. Biophys J 1995, 68(6):2588–2600.

21. Brakenhoff GJ, Visscher K, Gijsbers EJ: Fluorescence bleach rate imaging. J Microsc 1994, 175:154–161.

22. Young RM, Arnette K, Roess DA, Barisas BG: Quantitation of fluorescence energy transfer between cell surface proteins via fluorescence donor photobleaching kinetics. Biophys J 1994, 67:881–888.

23. Benson DM, Bryan J, Plant AL, Gotto AMJ, Smith LC: Digital imaging fluorescence microscopy: spatial heterogeneity of photobleaching rate constants in individual cells. J Cell Biol 1985, 100:1309–1323.

24. Van Oostveldt P, Verhaegen F, Messens K: Heterogeneous photobleaching in confocal microscopy caused by differences in refractive index and excitation mode. Cytometry 1998, 32(2):137–146.

25. Zwier JM, van Rooij GJ, Hofstraat JW, Brakenhoff GJ: Image calibration in fluorescence microscopy. J Microsc 2004, 216(1):15–24.

26. Schmid PJ: Dynamic mode decomposition of numerical and experimental data. J Fluid Mech 2010, 656:5–28.

27. Brunton SL, Kutz JN: Data-driven science and engineering: Machine learning, dynamical systems, and control. Cambridge: Cambrudge University Press; 2019.

28. Bi C, Yuan Y, Zhang JW, Shi Y, Xiang Y, Wang Y, Zhang RH: Dynamic Mode Decomposition Based Video Shot Detection. IEEE Access 2018, 6:21397–21407.

29. Kutz JN, Fu X, Brunton SL, Erichson NB: Multi-resolution dynamic mode decomposition for foreground/background separation and object tracking. IEEE International Conference on Computer Vision Workshop 2016.

30. Tirunagari S, Poh N, Wells K, Bober M, Gorden I, Windridge D: Functional Segmentation through Dynamic Mode Decomposition: Automatic Quantification of Kidney Function in DCE-MRI Images. arXivorg 2019.

31. Casorso J, Kong X, Chi W, Van De Ville D, Yeo BTT, Liegeois R: Dynamic mode decomposition of resting-state and task fMRI. Neuroimage 2019, 194:42–54.

32. Li J, Brown G, Ailion M, Lee S, Thomas JH: NCR-1 and NCR-2, the C. elegans homologs of the human Niemann-Pick type C1 disease protein, function upstream of DAF-9 in the dauer formation pathways. Development 2004, 131(22):5741–5752.

33. Lee HJ, Zhang W, Zhang D, Yang Y, Liu B, Barker EL, Buhman KK, Slipchenko LV, Dai M, Cheng JX: Assessing cholesterol storage in live cells and C. elegans by stimulated Raman scattering imaging of phenyl-Diyne cholesterol. Sci Rep 2015, 5:7930.

34. Maxfield FR, McGraw, T.E.: Endocytic recycling. Nat Rev Mol Cell Biol 2004, 5:121–132.

35. Hao M, Maxfield FR: Characterization of rapid membrane internalization and recycling. J Biol Chem 2000, 275:15279–15286.

36. Wüstner D, Solanko LM, Sokol E, Lund FW, Garvik O, Li Z, Bittman R, Korte T, Herrmann A: Quantitative Assessment of Sterol Traffic in Living Cells by Dual Labeling with Dehydroergosterol and BODIPY-cholesterol. Chem Phys Lipids 2011, 164(3):221–235.

37. Mayor S, Presley JF, Maxfield FR: Sorting of membrane components from endosomes and subsequent recycling to the cell surface occurs by a bulk flow process. J Cell Biol 1993, 121:1257–1269.

38. Presley JF, Mayor S, Dunn KW, Johnson LS, McGraw TE, Maxfield FR: he End2 mutation in CHO cells slows the exit of transferrin receptors from the recycling compartment but bulk membrane recycling is unaffected. J Cell Biol 1993, 122(6):1231–1241.

39. Presley JF, Mayor S, McGraw TE, Dunn KW, Maxfield FR: Bafilomycin A1 treatment retards transferrin receptor recycling more than bulk membrane recycling. J Biol Chem 1997, 272(21):13929–13936.

40. Demchenko AP: Photobleaching of organic fluorophores: quantitative characterization, mechanisms, protection. Methods Appl Fluoresc 2020, 8(2):022001.

41. Bernas T, Zaresbski M, Cook RR, Dobrucki JW: Minimizing photobleaching during confocal microscopy of fluorescent probes bound to chromatin: role of anoxia and photon flux. J Microscopy 2004, 215(3):281–296.

42. Modzel M, Solanko KA, Szomek M, Hansen SK, Dupont A, Nabo LJ, Kongsted J, Wüstner D: Live-cell imaging of new polyene sterols for improved analysis of intracellular cholesterol transport. Journal of microscopy 2018.

43. Isselstein M, Zhang L, Glembockyte V, Brix O, Cosa G, Tinnefeld P, Cordes T: Self-Healing Dyes-Keeping the Promise? The journal of physical chemistry letters 2020, 11(11):4462–4480.

44. Ronnlund D, Xu L, Perols A, Gad AK, Eriksson Karlstrom A, Auer G, Widengren J: Multicolor fluorescence nanoscopy by photobleaching: concept, verification, and its application to resolve selective storage of proteins in platelets. ACS Nano 2014, 8(5):4358–4365.

45. Lingvay M, Akhtar P, Sebok-Nagy K, Pali T, Lambrev PH: Photobleaching of Chlorophyll in Light-Harvesting Complex II Increases in Lipid Environment. Front Plant Sci 2020, 11:849.

46. Szabo G, Pine PS, Weaver JL, Kasari M, Aszalos A: Epitope mapping by photobleaching fluorescence resonance energy-transfer measurements using a laser scanning microscope system. Biophys J 1992, 61:661–670.

47. Tramier M, Zahid M, Mevel JC, Masse MJ, Coppey-Moisan M: Sensitivity of CFP/YFP and GFP/mCherry pairs to donor photobleaching on FRET determination by fluorescence lifetime imaging microscopy in living cells. Microsc Res Tech 2006, 69(11):933–939.

48. Liu X, Pomorski TG, Liesche J: Non-invasive Quantification of Cell Wall Porosity by Fluorescence Quenching Microscopy. Bio Protoc 2019, 9(16):e3344.

49. Lehrer SS: Solute perturbation of protein fluorescence. The quenching of the tryptophyl fluorescence of model compounds and of lysozyme by iodide ion. Biochemistry 1971, 10(17):3254–3263.

50. Hale JE, Schroeder F: Asymmetric transbilayer distribution of sterol across plasma membranes determined by fluorescence quenching of dehydroergosterol. Eur J Biochem 1982, 122(3):649–661.

51. Mondal M, Mesmin B, Mukherjee S, Maxfield FR: Sterols are mainly in the cytoplasmic leaflet of the plasma membrane and the endocytic recycling compartment in CHO cells. Mol Biol Cell 2009, 20(2):581–588.

52. Solanko LM, Sullivan DP, Sere YY, Szomek M, Lunding A, Solanko KA, Pizovic A, Stanchev LD, Pomorski TG, Menon AK et al: Ergosterol is mainly located in the cytoplasmic leaflet of the yeast plasma membrane. Traffic 2018, 19(3):198–214.

53. White JC, Stryer L: Photostability studies of phycobiliprotein fluorescent labels. Anal Biochem 1987, 161:442–452.

54. Georgakoudi I, Foster TH: Singlet Oxygen-Versus Nonsinglet Oxygen-Mediated Mechanisms of Sensitizer Photobleaching and Their Effects on Photodynamic Dosimetry. Photochem Photobiol 1998, 67(6):612–625.

55. Albro PW, Bilski P, Corbett JT, Schroeder JL, Chignell CF: Photochemical reactions and phototoxicity of sterols: novel self-perpetuating mechanisms for lipid photooxidation. Photochemistry and photobiology 1997, 66(3):316–325.

56. Spikes JD: Quantum yields and kinetics of the photobleaching of hematoporphyrin, Photofrin II, tetra(4-sulfonatophenyl)-porphine and uroporphyrin. Photochemistry and photobiology 1992, 55(6):797–808.

57. Yamashiro DJ, Tycko, B., Fluss, S.R., and F.R. Maxfield.: Segregation of transferrin to a mildly acidic (pH 6.5) para-Golgi compartment in the recycling pathway. Cell 1984, 37:789–800.

58. Wüstner D, Mukherjee S, Maxfield FR, Müller P, Hermann A: Vesicular and nonvesicular transport of phosphatidylcholine in polarized HepG2 cells. Traffic 2001, 2:277–296.

59. Iversen TG, Skretting G, van Deurs B, Sandvig K: Clathrin-coated pits with long, dynamin-wrapped necks upon expression of a clathrin antisense RNA. Proc Natl Acad Sci U S A 2003, 100(9):5175–5180.

60. Wüstner D, Solanko LM, Lund FW, Sage D, Schroll JA, Lomholt MA: Quantitative fluorescence loss in photobleaching for analysis of protein transport and aggregation. BMC Bioinformatics 2012, 13:296.

61. Demo N, Tezzele M, Rozza G: PyDMD: Python Dynamic Mode Decomposition. J Open Source Software 2018, 3(22):530.

62. PixBleach: Pixelwise analysis of bleach rate in time-lapse images. A plugin to ImageJ. [http://bigwww.epfl.ch/algorithms/pixbleach/]

