## Supplemental_data for "Image segmentation and separation of spectrally similar dyes in fluorescence microscopy by dynamic mode decomposition of photobleaching kinetics"

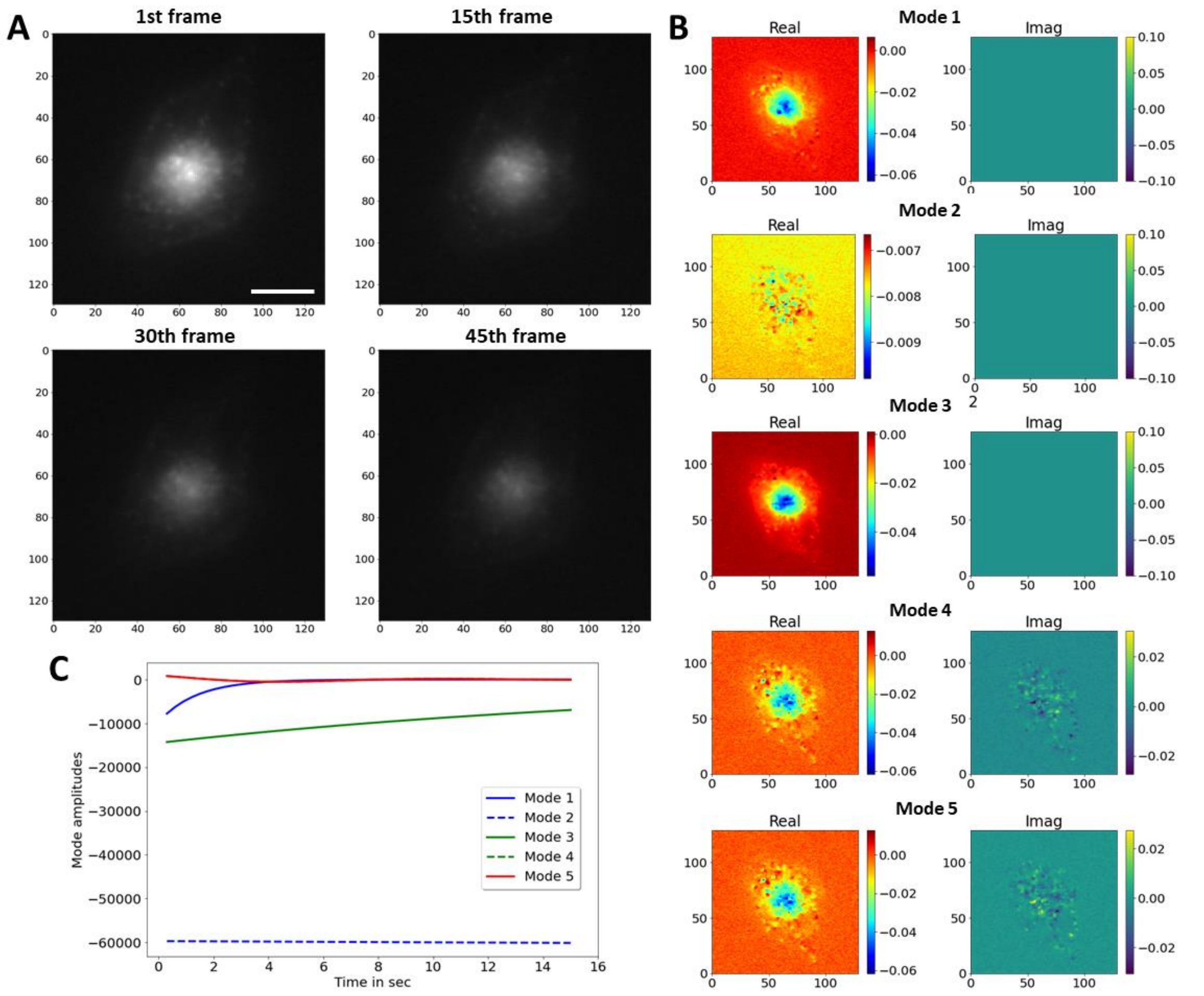


**Additional File 1: Figure S1. Dynamic mode decomposition of image stacks containing Alexa488-Tf labeled cells.** BHK cells were labeled with 20 µg/ml Alexa488-Tf for 30 min, washed with buffer medium and imaged on a wide field fluorescence microscope. A, selected frames of an image stack acquired with 0.3 sec acquisition time and without pause. Images are identically scaled; bar 10 µm. B, C, DMD of this image stack using a rank-5 approximation to the full transfer matrix. B, mode weights and C, mode amplitudes as function of time.

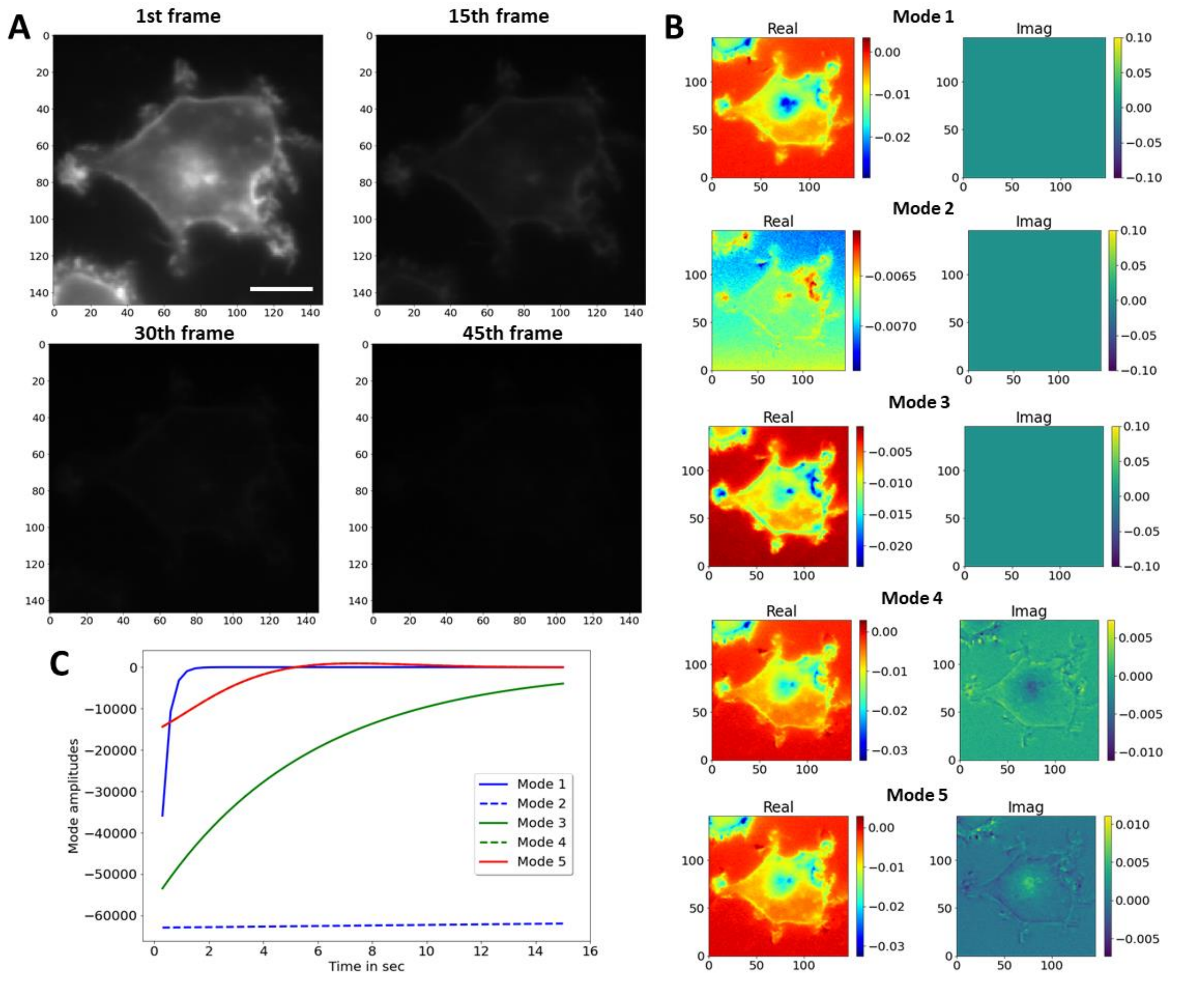


**Additional File 2: Figure S2. Dynamic mode decomposition of image stacks containing C6-NBD-SM labeled cells.** BHK cells were labeled with 4 µM C6-NBD-SM for 30 min, washed with buffer medium and imaged on a wide field fluorescence microscope. A, selected frames of an image stack acquired with 0.3 sec acquisition time and without pause. Images are identically scaled; bar 10 µm. B, C, DMD of this image stack using a rank-5 approximation to the full transfer matrix. B, mode weights and C, mode amplitudes as function of time.

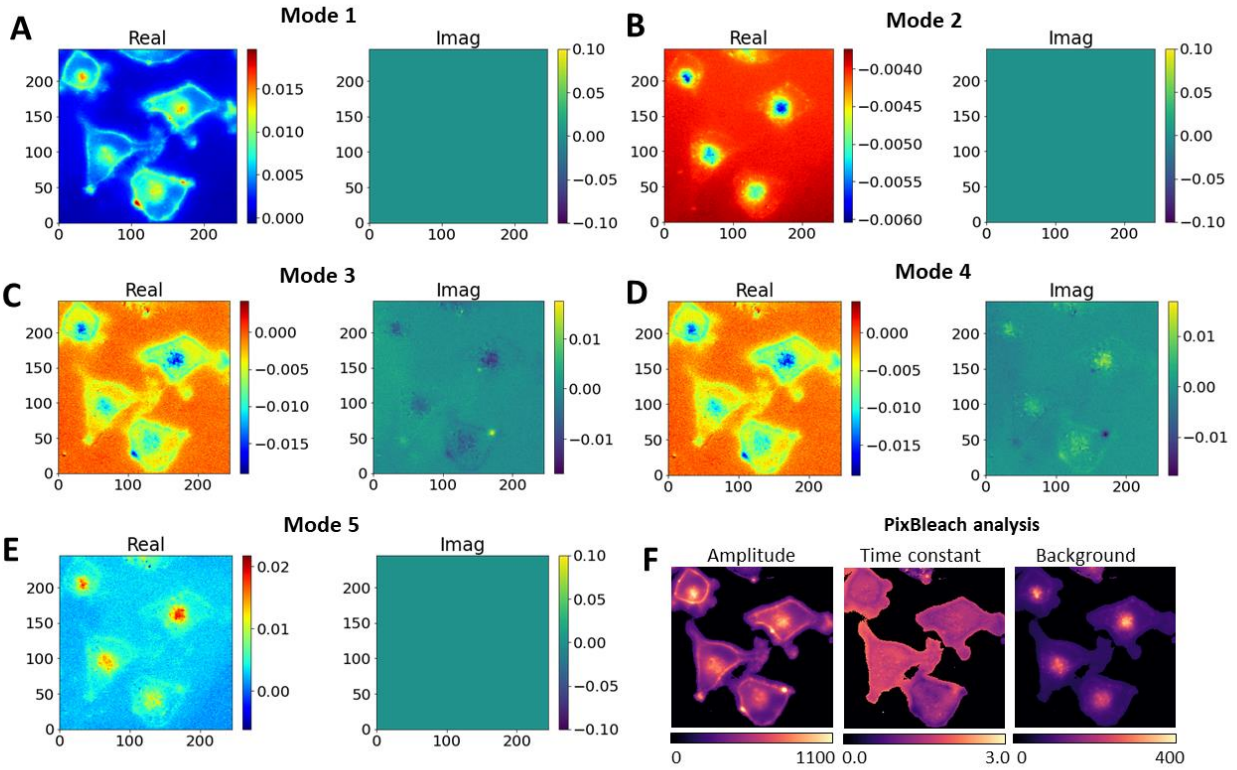


**Additional File 3: Figure S3. Mode weights for DMD of image stacks of BHK cells labeled with C6-NBD-SM and Alexa488-Tf.** BHK cells were labeled with 4 µM C6-NBD-SM and with 20 µg/ml Alexa488-Tf, both emitting in green, as described in Materials and Methods. BHK cells were labeled with 4 µM C6-NBD-SM for 30 min, washed with buffer medium and imaged on a wide field fluorescence microscope. Mode weights for DMD of rank 5 of this data are shown. The real part of mode weights is shown in left panels ('Real'), while the imaginary parts are shown in right panels ('Imag'). F, bleach rate fitting using a stretched exponential function with bleaching amplitudes (right panel), time constant (middle panel) and background term (left panel).

**Additional file 4: Simulated bleach stack.** Raw data set 1 used in the analysis shown in Figures 1 to 3. Photobleaching was simulated using single-exponential decay functions as described in Materials and Methods.

**Additional file 5: Experimental bleach stack of C. elegans labeled with DHE.** Raw data set 2 used in the analysis shown in Figure 4 to 7. C. elegans was labeled with DHE, and images were acquired with 0.5 sec acquisition time and without pause as described in Materials and Methods.

**Additional file 6: Experimental bleach stack of BHKasc cells double-labeled with C6-NBD-SM and Alexa488-Tf.** Raw data set 3 used in the analysis shown in Figure 8 and S3. BHKasc cells were labeled with C6-NBD-SM and Alexa488-TF, and images were acquired with 0.3 sec acquisition time and without pause as described in Materials and Methods.

**Additional file 7: Experimental bleach stack of BHKasc cells labeled with Alexa488-Tf.** Raw data set 4 used in the analysis shown in Figure S1. BHKasc cells were labeled with Alexa488-TF, and images were acquired with 0.3 sec acquisition time and without pause as described in Materials and Methods.

**Additional file 8: Experimental bleach stack of BHKasc cells labeled with C6-NBD-SM.** Raw data set 5 used in the analysis shown in Figure S2. BHKasc cells were labeled with C6-NBD-SM, and images were acquired with 0.3 sec acquisition time and without pause as described in Materials and Methods.
